# A combination of Interferon Stimulated Genes is more effective than IFNα and IFNβ in reducing HIV-1 replication in human cervicovaginal tissues

**DOI:** 10.1101/610964

**Authors:** Christiane Rollenhagen, Jiang Gui, Gustavo F. Doncel, Susana N. Asin

## Abstract

Enhancing antiviral responses while controlling immune cell activation is an attractive strategy to reduce HIV-1 replication in the cervicovaginal mucosae, a primary site of heterosexual transmission. Interferon alpha and beta (IFNα/β) signaling up-regulates expression of inflammatory factors and Interferon-Stimulated Genes (ISGs). The simultaneous induction of both IFNs by pathogen-bound molecular pattern recognition receptors and the paucity of data on the anti-HIV-1 efficacy of a combination of these antiviral factors or their downstream targets in human experimental models taking into account mucosal and submucosal cell populations, motivated us to determine whether combined IFNα/β or ISGs could decrease HIV-1 replication in cervicovaginal tissues.

IFNα/β reduced HIV-1 p24 release. This reduction was associated with upregulation of expression of a subset of ISGs, the type I IFN receptor and interferon regulatory factor seven. IFNα/β also enhanced immune cell activation. In contrast, when added directly to CV tissues, a combination of ISGs was more effective than IFNα/β in reducing HIV-1 p24 release. The ISG combination demonstrated early kinetics and a more robust reduction in HIV-1 p24 release. Opposite to IFNα/β, the combination of ISGs did not induce immune cell activation.

IFNα/β-induced ISGs provide novel mucosal therapeutic targets with a greater capacity to reduce HIV-1 compared to IFNα/β, without inducing immune cell activation.

## Introduction

Viral pathogen sensing by molecular pattern recognition receptors (PRRs), activates signaling pathways to produce type I Interferons (IFNs) [1-4]. The IFNα and β stimulated host innate immune response to viral pathogens is driven by IFN regulatory factors (IRFs), mainly IRF3 and IRF7 [4-9]. IRF3 is constitutively expressed and undergoes phosphorylation and nuclear translocation following viral infection, whereas IRF7 is mostly induced by HIV-1 activation of PRRs. Upon phosphorylation IRF3 forms homo- or heterodimers with IRF7, which translocate to the nucleus and drive expression of interferon-stimulated genes (ISGs) and additional genes that modulate cell cycle progression and apoptosis [10].

The antiviral effects of IFNα or β administration in animal models have been studied and generated contradicting results. Intramuscular applications of pegylated recombinant (r)IFNα2 to female *Rhesus macaques* delayed but did not prevent infection by rectally inoculated Simian Immunodeficiency Virus (SIV) in treated compared to placebo control animals [11]. This transient protection was associated with enhanced expression of ISGs. Consistent with this finding, a type I IFN receptor (IFNR) antagonist enhanced plasma viral load and delayed ISG expression in the peripheral blood of antagonist treated compared to untreated control animals [11]. A more recent study demonstrated that vaginal administrations of a human(h) rIFNβ protected 9 of 15 treated *R. macaques* from Simian Human Immunodeficiency Virus (SHIV) infection [12]. The protective effect was associated with the induction of ISGs in hrIFNβ treated compared to untreated animals [12]. Likewise, plasmids encoding subtypes of IFNα or IFNβ reduced HIV-1 viral load in severe combined immunodeficiency mice grafted with human peripheral blood (hu)-PBL mice [13,14]. Antiviral activity differed among IFNα sub-types. Supporting these findings, a lack of effect or even a stimulation of viral replication by rIFNα has also been reported [15,16]. IFNα enhanced both HIV-1 plasma viral load and T cell activation in the hu-bone marrow-liver-thymus (BLT) mice [16].

To our knowledge no studies have addressed the effect of combined IFNα and β in human cervicovaginal (CV) tissues. Human experimental models have been limited to primary cells and cell lines. Studies in primary macrophages and CD4^+^ T cells demonstrated that IFNα supplementation blocked early and late steps of the HIV-1 life cycle, whereas a significant reduction in HIV-1 replication was reported in human CD4^+^ T cells expressing IFNβ [17-19]. Given the simultaneous induction of IFNα and β expression by sexually transmitted viruses, and the paucity of data in human experimental systems accounting for the interaction among mucosal and submucosal cell populations [20-22], we evaluated the impact of combined hrIFNα and β (hrIFNα/β) on HIV-1 infection in CV tissues. HrIFNα/β reduced HIV-1 p24 release in CV tissue supernatants. Lower levels of viral replication were associated with an unchanged expression of the HIV-1 receptor CD4, and enhanced expression of the HIV-1 co-receptor CCR5 and immune cell activation marker CD38. HrIFNα/β up-regulated expression of the IFNα and β receptor Subunit 2 (IFNAR2), the cellular transcription factor IRF7, ISG15, bone marrow stromal cell antigen (BST)2, myxovirus resistance protein (MX)2 and interferon induced transmembrane protein (IFITM)1.

These findings suggest that the reduction in HIV-1 p24 release by hrIFNα/β was the result of the antiviral effect of ISGs counterbalancing mucosal immune cell activation. Supporting our postulate, we demonstrated that exogenous supplementation with hrISG15 and hrBST2, two downstream targets induced by hrIFNα/β resulted in an earlier and more significant reduction in HIV-1 p24 release compared to hrIFNα/β. In contrast to hrIFNα/β, the combination of these ISGs did not induce immune cell activation.

Our original findings underscore the potential of using hrISGs rather than hrIFNs as therapeutic targets to more effectively reduce HIV-1 replication without inducing immune cell activation in the CV mucosae.

## Materials and Methods

### Tissue samples

CV tissues were procured from HIV-1 sero-negative women who were undergoing hysterectomy at Dartmouth-Hitchcock Medical Center (DHMC) for benign medical conditions. This study received a Human Subjects Research no engaged determination by the Committee for the Protection of Human Subjects at Dartmouth College, and an exempt Category #5 determination by the Northern New England Research Consortium VA Medical Centers Institutional Review Board; based on the fact that CV tissue specimens were collected for non-research purposes and investigators did not have access to any Personally Identifiable or Protected Health Information. Non-polarized CV tissue cultures were established in 48-well plates as described [8, 23-25]. Using our culture conditions, tissue explants are maintained for up to 21 days without decrease in viability, as determined by Lactate Dehydrogenase viability assay (Cytotoxicity Detection Kit, Roche, Indianapolis, IN).

### HIV-1 infection

The R5-tropic HIV-1_BaL_ stock was generated in human peripheral blood mononuclear cells. Tissues were infected with 10^4^ 50% Tissue Culture Infectious Dose (TCID50)/ml of cell-free HIV-1_BaL_. After overnight incubation at 37°C, tissues were washed to remove residual input virus, and cultured for up to 21 days in Leibowitz (L) 15 medium supplemented with 10% heat inactivated fetal bovine serum (Hyclone, Logan, UT), 2 mM glutamine (GIBCO, Grand Island, NY), 50 unit/ml penicillin and 50 μg/ml streptomycin (complete L15, GIBCO, Grand Island, NY). Culture supernatants were collected after the final wash (day 0), and on days 4, 7, 11, 14, 18 and 21 after infection. On each day, one half of the culture supernatant was removed, and replenished with an equivalent volume of fresh media. We evaluated supernatants from days 0, 11, 14 and 21 for HIV-1 p24 antigen levels by ELISA (Perkin Elmer, Boston, MA).

### Exogenous addition of human recombinant (hr)proteins

HrIFNα (a hybrid of subtypes one and two at 10^5^ units/ml, Biotechne, Minneapolis, MN) in combination with hrIFNβ at 2 ng/ml (R&D Systems, Minneapolis, MN) or when indicated hrISG15 at 0.115 μg/ml (Biotechne, Minneapolis, MN) alone or in combination with a hrBST2 peptide at 0.1 μg/ml (iba, Goettingen, Germany) were added to CV tissues before, during and after the infection. The hrIFNα and IFNβ concentrations were selected in titration experiments testing three IFNα (10^3^, 10^4^ and 10^5^ units /ml) and three IFNβ (2, 0.2 and 0.02 ng/ml) concentrations. These concentrations are in the range of those evaluated in primary human CD4^+^ T cells, macrophages and cell lines (ref). Likewise, the hrISG15 and hrBST2 concentrations were selected in titration experiments testing ISG15 at 1.15, 0.115 and 0.0115 μg/ml and BST2 at 1, 0.1 and 0.01 μg/ml. The selected combinations of hrproteins or peptide demonstrated the highest reduction in HIV-1 p24 release without inducing tissue toxicity as indicated with a LDH assay (Pierce LDH Cytotoxicity Assay, Thermo Scientific, Rockford, IL).

### Nucleic acid isolation

RNA was isolated from CV tissues on days 5 and 7 after infection. Tissues were homogenized and lysate supernatants were subjected to RNA isolation using the RNeasy-Plus kit (Qiagen, Valencia, CA).

### Gene Transcription

One μg of total tissue RNA was reverse transcribed using Superscript III reverse transcriptase (Invitrogen, Carisbad, CA). One µl of the resulting cDNA conversion was evaluated for gene expression by real-time PCR using SsoAdvUniver SYBR GRN SMX 2500 (BioRad, Hercules, California). All transcription values were normalized to endogenous human Glyceryl aldehyde 3-Phosphate (GAPDH). Primer sequences are described in Table 1.

**Table 1.**
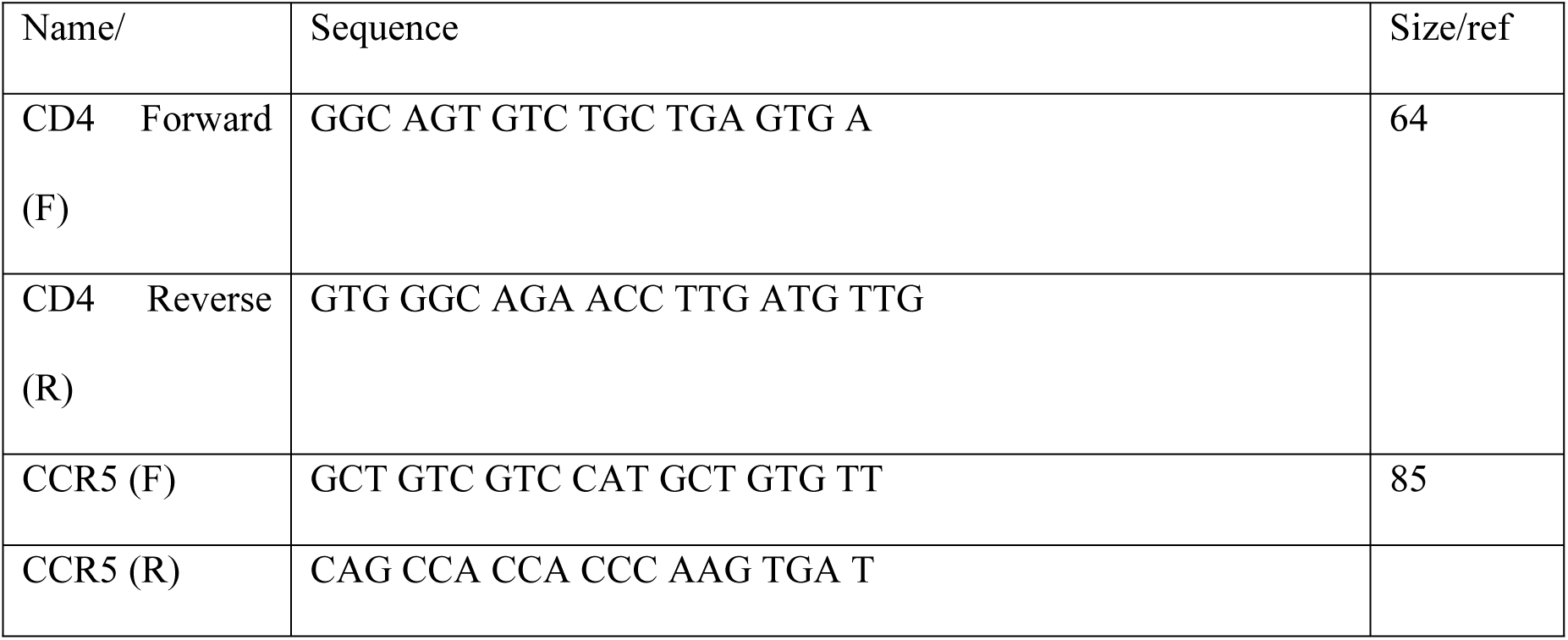
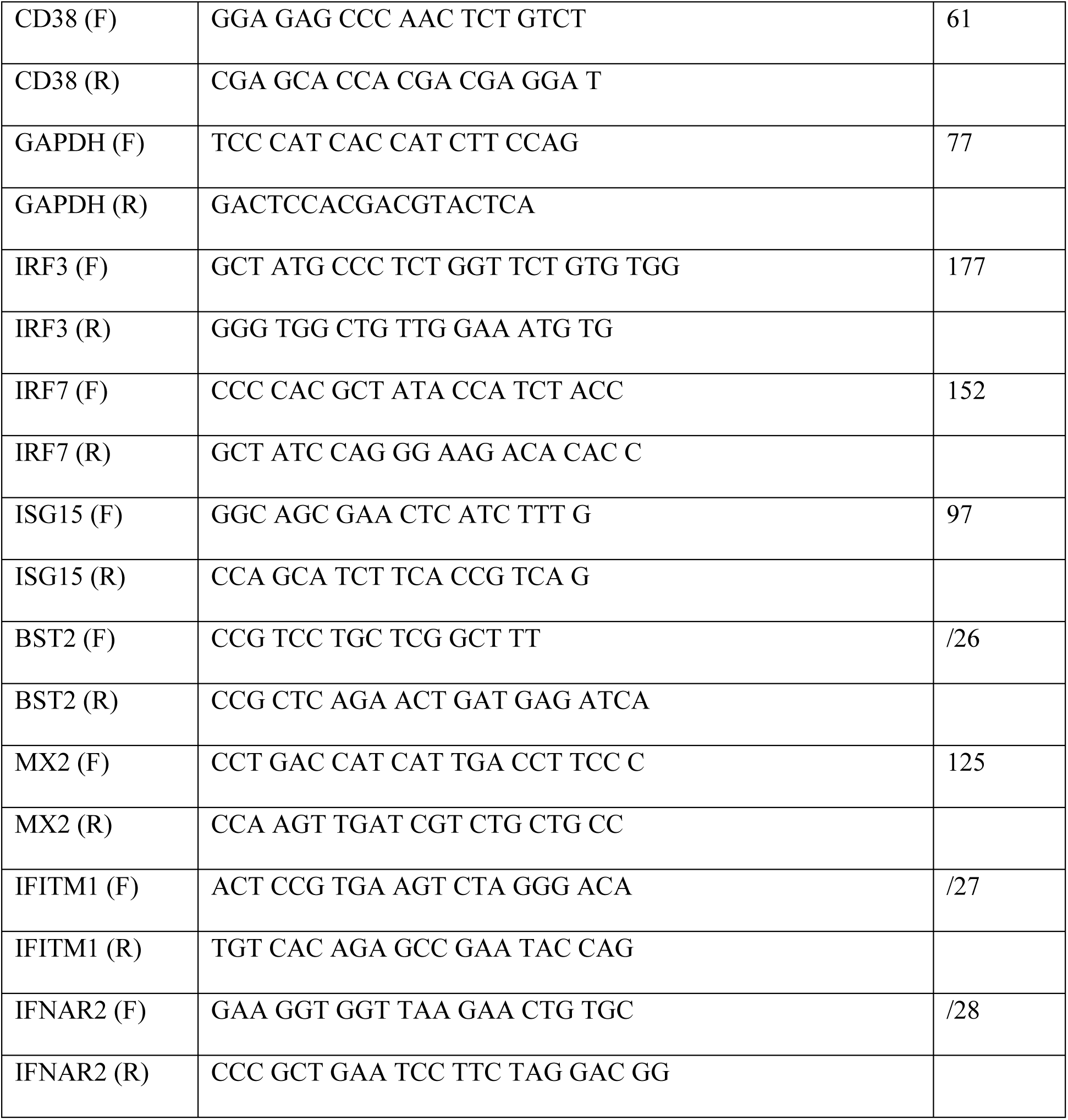
Sequence of primer used to amplify target genes

### Statistical analyses

Datasets containing two groups (untreated and hrproteins/peptide treated tissues) at days 11, 14 and 21 of infection were not normally distributed and were analyzed by Wilcoxon rank sum non-parametric test. We used GraphPad Prism version 6.01 (GraphPad, SanDiego, CA, USA) for graphs. P values of <0.05 were considered significant. Based on our power calculation, we need to analyze data from at least 15 tissues to achieve a 90% power for a type I error of 5 % and an effect size of 90%.

## Results

### Human recombinant (hr) IFNαβ decreases HIV-1 replication in CV tissues

Neither IFNα nor β alone reduced HIV-1 p24 release in CV tissues (data not shown). Our published data suggest that the PRR ligand Polyinosinic:polycytidylic acid [Poly (I:C)] reduces HIV-1 replication in CV tissues by enhancing IFNα and β expression [8]. To test the efficacy of these antiviral factors at decreasing HIV-1 infection, we treated CV tissues with a combination of human recombinant IFNα and β proteins (hrIFNα/β). In these experiments, CV tissues were established as described [8,23-25], and left untreated or treated with hrIFNα/β (IFNα at10^5^ units/ml and IFNβ at 2 ng/ml) prior to infection with R5-tropic HIV-1_BaL_. After overnight incubation, tissues were extensively washed to remove residual input virus and cultured for 21 days. After washing (Day 0), hrIFNα/β was added back to designated tissues and replenished every three days. Untreated tissues were included as controls. Tissue culture supernatants were evaluated for HIV-1 p24 levels on days 11 and 21 of infection. P24 is an HIV-1 antigen widely used as a surrogate for HIV-1 release. In general, p24 levels peak at day 11 and we terminate experiments on day 21 [23, 24]. Since residual input virus is mainly released through day 7 of infection, p24 levels at days 11 and 21 mostly reflect the production of newly synthesized virus [25]. Results from these experiments demonstrated decreased HIV-1 p24 levels in supernatants from tissues treated with hrIFNα/β compared to untreated control tissues at day 21 of infection (Fig. 1).

**Figure 1.**
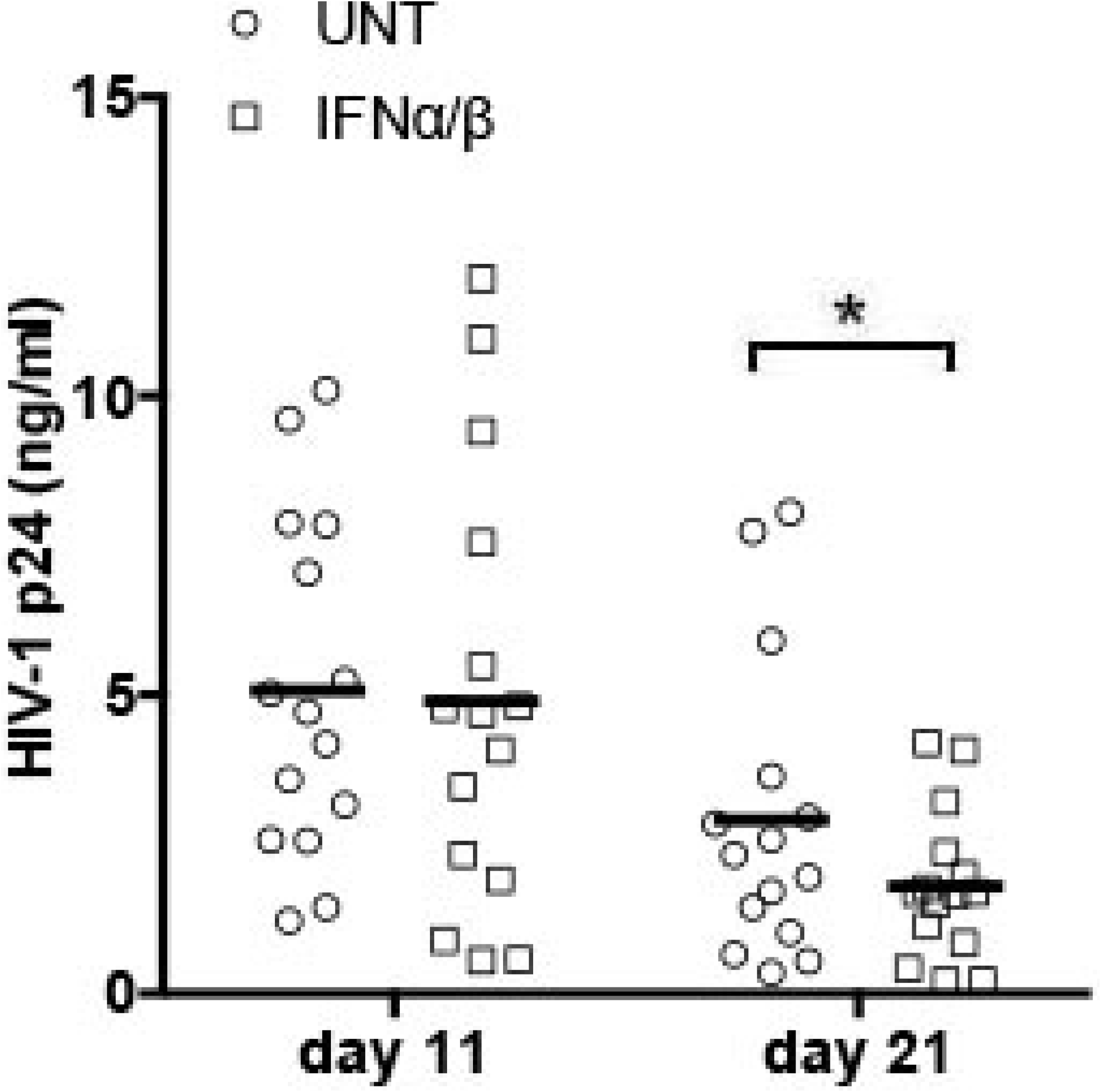
HrIFNα/β decreases HIV-1 replication in cervicovaginal (CV) tissues. HIV-1 p24 levels (ng/ml) in HIV-1 infected CV tissues left untreated or treated with 10^5^ units/ml of hrIFNα in combination with 2 µg/ml of hrIFNβ were measured after washing the residual input virus (day 0), and again on days 11 and 21 after infection. Mean values of triplicates from 15 individual donors are shown. HIV-1 p24 levels on day 0 were below the limit of detection of the assay and are not depicted in the figure. * p< 0.05 comparing untreated (UNT) to IFNα/β treated CV tissues.

The reduction in HIV-1 p24 release was not a result of hrIFNα/β mediated tissue toxicity as indicated by lactate dehydrogenase (LDH) levels in tissue culture supernatants (data not shown).

### HrIFNα/β enhances immune cell activation in CV tissues

As HIV-1 mainly replicates in activated CD4^+^ T cells [29], we tested whether the decrease in HIV-1 p24 release by hrIFNα**/**β was associated with a lack of immune cell activation. We compared expression levels of CD4, CCR5 and CD38, between hrIFNα**/**β treated and untreated control tissues on days 5 and 7 of infection. We selected these time points to evaluate gene transcription following activation but before immune cell depletion due to the progression of HIV-1 infection [23]. Even though hrIFNα/β did not alter CD4 expression (Fig. 2A), we found enhanced CCR5 and CD38 expression in hrIFNα/β treated compared to untreated tissues at day 7 of infection (Fig. 2 B and C)

**Figure 2.**
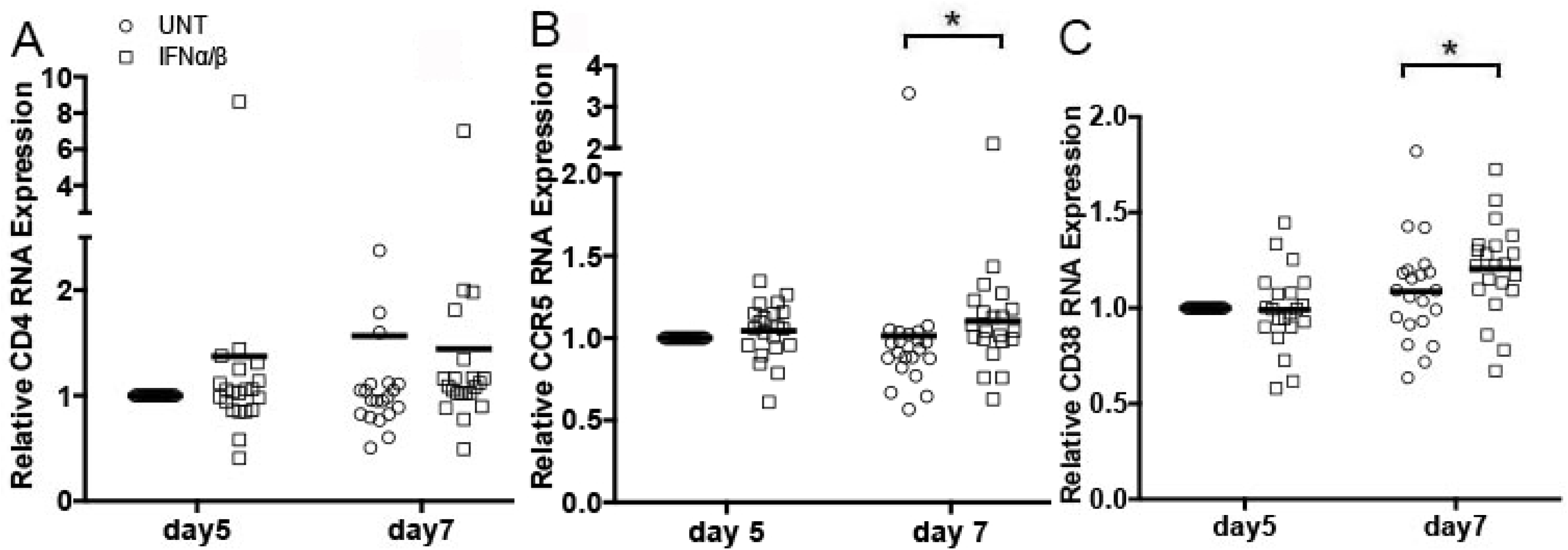
HrIFNα/β induces immune cell activation. RNA expression levels of the HIV-1 receptor CD4 (A), co-receptor CCR5 (B) and the immune cell activation marker CD38 (C) in CV tissues left untreated or treated with 10^5^ units/ml of hrIFNα in combination with 2 µg/ml of hrIFNβ were quantified by RT-PCR on days 5 and 7 after infection. All data were normalized to Glyceraldehyde 3-phosphate (GAPDH). For each gene, day 5 values in untreated control tissues were set to 1. Day 5 values in IFNα/β treated tissues or day 7 values in untreated and IFNα/β treated tissues were normalized to day 5 values in untreated tissues. Mean values of triplicates from 21 individual donors are shown. * p< 0.05 comparing untreated (UNT) to IFNα/β treated CV tissues.

### A combination of hrIFNα/β enhances IFNAR2 expression

HrIFNα/β signals through dimerization of the two IFN α and β receptor (IFNAR) subunits 1 and 2. To address whether the hrIFNα/β mediated reduction in HIV-1 replication was associated with the stimulation of an innate immune anti-viral signaling pathway, we compared RNA expression of the IFNAR2 between hrIFNα/β treated and untreated tissues on days 5 and 7 of infection. We selected IFNAR2 because of the reported high binding affinity of hrIFNα/β to this sub-unit [30]. IFNAR2 expression was enhanced at day 5 of infection in hrIFNα/β treated compared to untreated control tissues (Fig. 3). This effect was sustained until day 7 but was not statistically significant at this time point.

**Figure 3.**
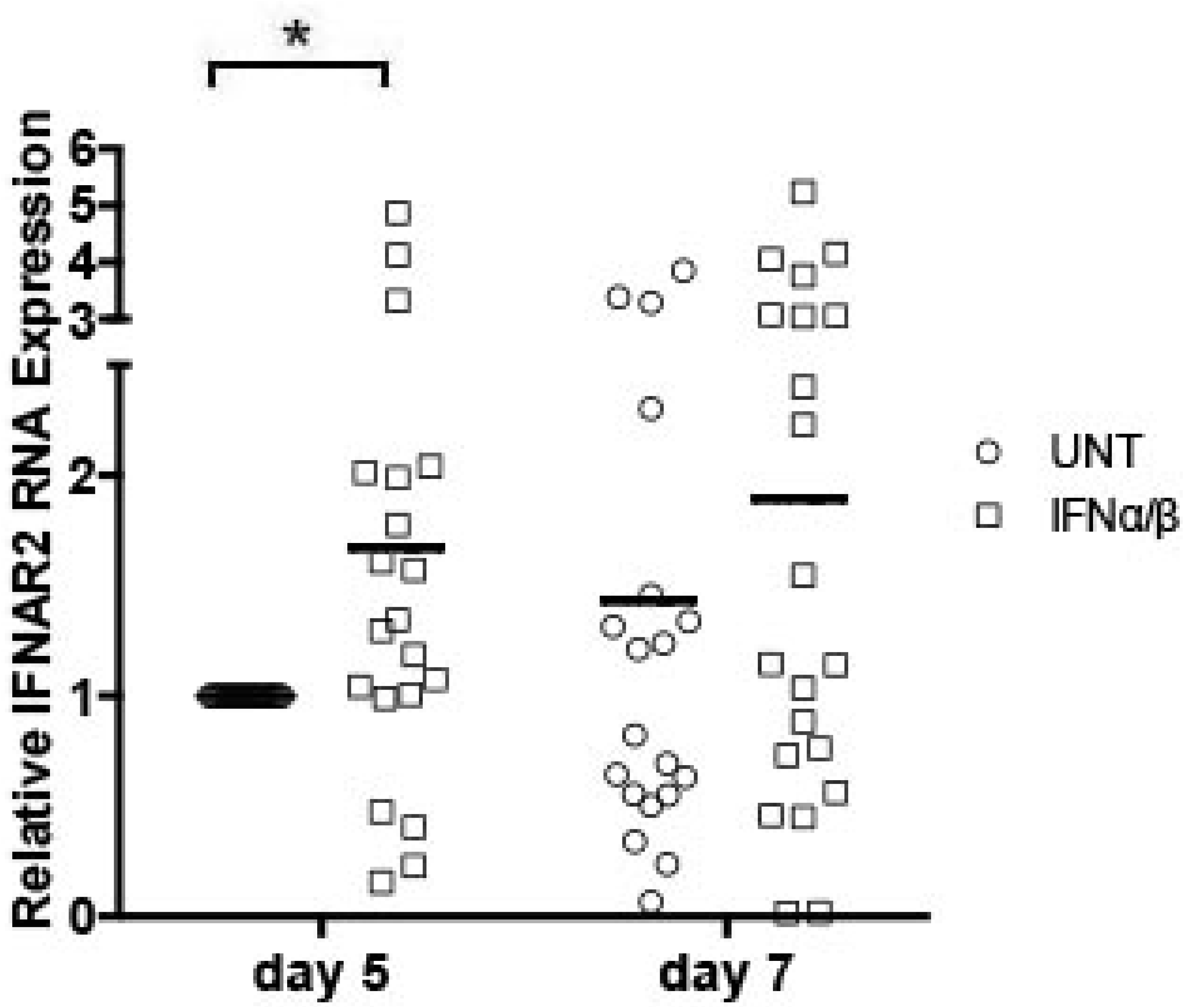
HrIFNα/β increases expression of the IFNα and β receptor Subunit 2 (IFNAR2). Levels of IFNAR2 in CV tissues left untreated or treated with 10^5^ units/ml of hrIFNα in combination with 2 µg/ml of hrIFNβ were quantified by RT-PCR on days 5 and 7 after infection. All data were normalized to GAPDH. Day 5 values in untreated tissues were set to 1. Day 5 values in IFNα/β treated tissues and day 7 values in untreated or IFNα/β treated tissues were normalized to day 5 values in untreated tissues. Mean values of triplicates from 21 CV tissues are shown. * p< 0.05 comparing untreated (UNT) to IFNα/β treated CV tissues.

### A combination of hrIFNα/β enhances Interferon Regulatory Factor (IRF)7 expression

Greater IFNAR2 RNA expression in hrIFNα/β treated compared to untreated tissues suggested the induction of an innate immune antiviral response likely mediated by Interferon Stimulated Genes (ISGs). Given that IRF3 and 7 are master regulators of ISG expression [9,10] with an essential role as antiviral effectors, we next compared expression of these IRFs between hrIFNα/β treated and untreated control tissues. HrIFNα/β enhanced IRF7 expression on day 7 of infection (Fig. 4B). In contrast, we found no differences in IRF3 expression levels between hrIFNα/β treated and untreated control tissues (Fig. 4A).

**Figure 4.**
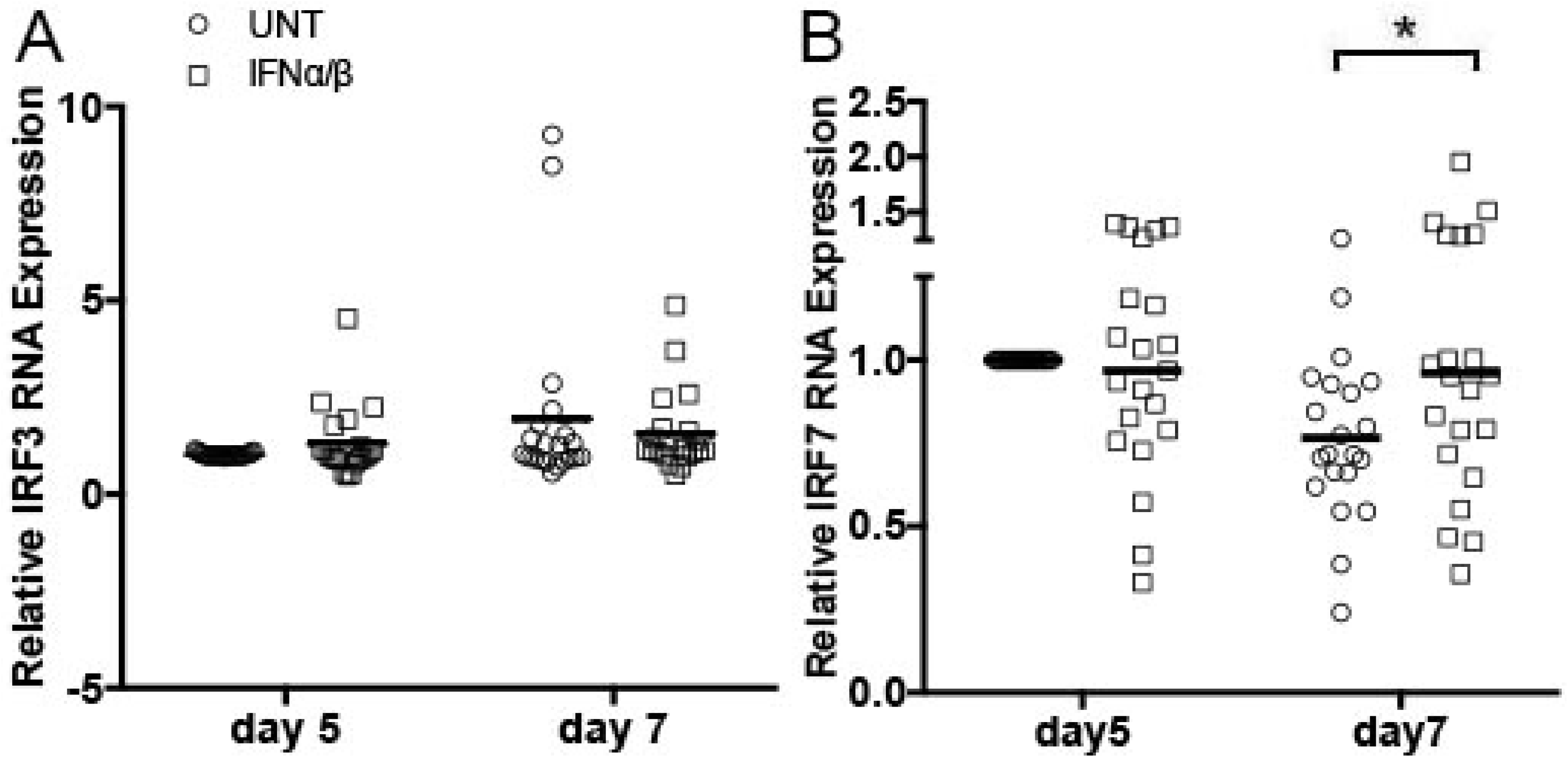
HrIFNα/β increases IRF7 expression. Levels of the cellular transcription factors IRF3 (A) and IRF7 (B) in CV tissues left untreated or treated with 10^5^ units/ml of hrIFNα in combination with 2 µg/ml of hrIFNβ were quantified by RT-PCR on days 5 and 7 after infection. All data were normalized to GAPDH. For each gene, day 5 values in untreated tissues were set to 1. Day 5 values in IFNα/β treated tissues and day 7 values in untreated or IFNα/β treated tissues were normalized to day 5 values in untreated tissues. Mean values of triplicates from 21 CV tissues are shown. * p< 0.05 comparing untreated (UNT) to IFNα/β treated CV tissues.

### A combination of hrIFNα/β enhances expression of ISGs

The enhanced expression of IRF7 in CV tissues treated with hrIFNα/β compared to untreated control tissues suggested the induction of an antiviral response mediated by ISGs. To test this hypothesis, we compared RNA levels of several ISGs including ISG15, BST2, MX2 and IFITM1 between hrIFNα/β treated and untreated control tissues on days 5 and 7 of infection. On day 7, we detected a significant up-regulation in ISG15, BST2 and MX2 expression by hrIFNα/β (Fig. 5A, B and C) whereas IFITM1 expression was enhanced at day 5 of infection (Fig. 5D).

**Figure 5.**
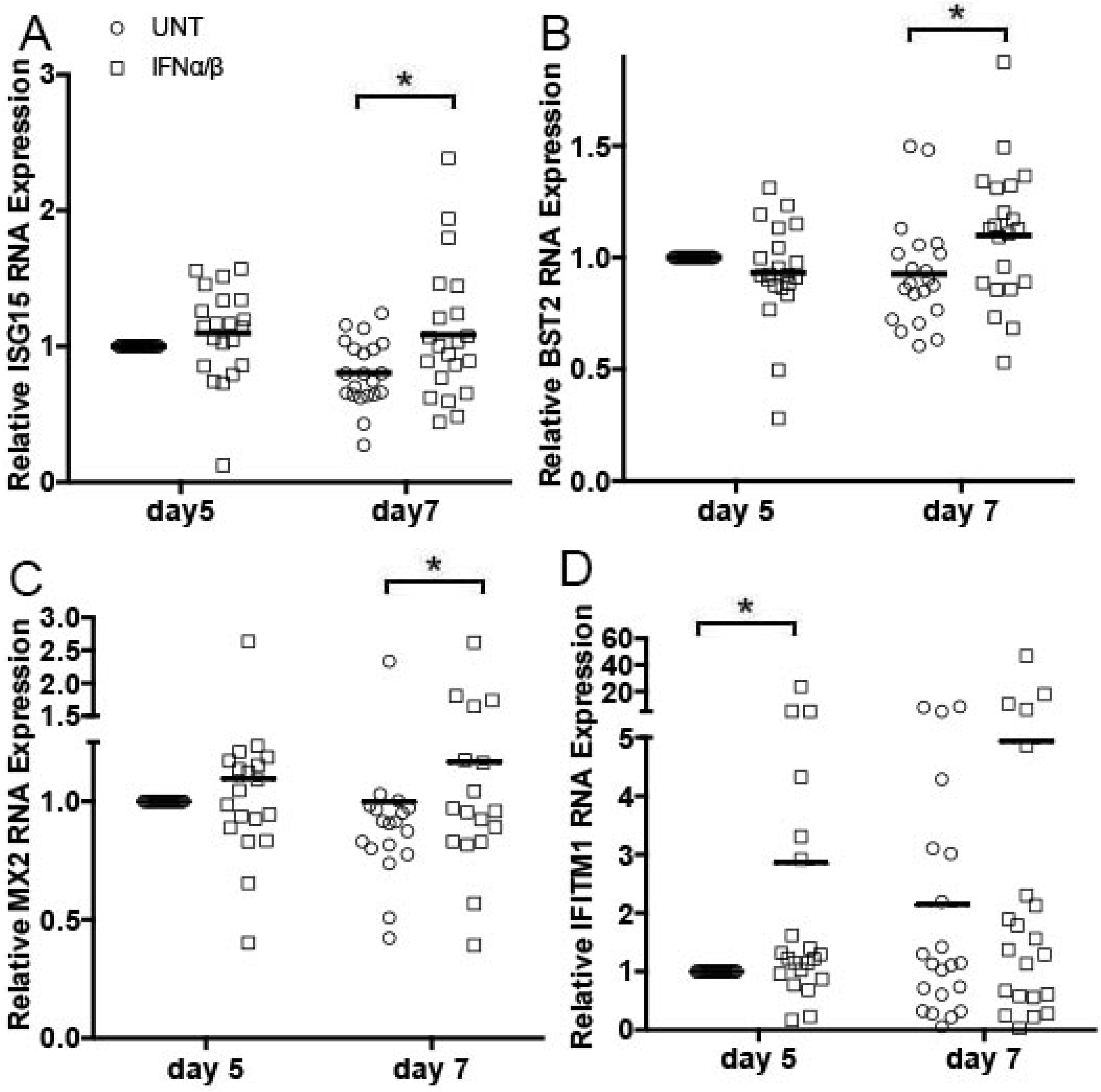
HrIFNα/β enhances ISG expression in cervicovaginal (CV) tissues. Levels of ISG15 (A), BST2 (B), MX2 (C) and IFITM1 (D) in CV tissues left untreated or treated with 10^5^ units/ml of hrIFNα in combination with 2 µg/ml of hrIFNβ, were quantified by RT-PCR on days 5 and 7 after infection. All data were normalized to GAPDH. For each gene, day 5 values in untreated tissues were set to 1. Day 5 values in IFNα/β treated tissues and day 7 values in untreated or IFNα/β treated tissues were normalized to day 5 values in untreated tissues. Mean values of triplicates from 21 CV tissues are shown. * p< 0.05 comparing untreated (UNT) to IFNα/β treated CV tissues.

### A Combination of ISGs is more effective than hrIFNα/β at reducing HIV-1 replication

HrIFNα/β reduced HIV-1 p24 release from CV tissues yet this decrease was observed on day 21 of infection. Given that hrIFNα/β induced expression of the well characterized antiviral effectors ISG15 and BST2 [31-34] in CV tissues, we postulated that these downstream targets will be more effective in reducing viral replication than hrIFNα/β supplementation. Furthermore, as both ISG15 and BST2 counteract viral release from infected cells by different mechanisms [31-34], we anticipated to detect a greater reduction in HIV-1 p24 release from tissues exposed to a combination of hrISG15 and a hrBST2 peptide (hrISG15/hrBST2) compared to those treated with single hrISG15. To test these hypotheses, we exposed CV tissues to hrISG15 (0.115 μg/ml) alone or in combination with a hrBST2 peptide (0.1 μg/ml) prior to infection with R5-tropic HIV-1_BaL_. HrISG15 and hrBST2 peptide concentrations were determined in titration experiments as described below (Methods). After infection overnight and washing of the residual input virus, hrISG15/hrBST2 were added back to designated tissues and replenished every three days. Untreated tissues and those treated with hrISG15 alone were included as controls. Results from these experiments revealed a significant reduction in HIV-1 p24 release at day 11 of infection in supernatants from CV tissues treated with combined hrISG15 and a hrBST2 peptide (hrISG15/BST2) compared to untreated control tissues. This reduction was even more pronounced at day 14 but was not sustained until day 21 of infection (Fig. 6). HrISG15 alone was not effective compared to the combination in reducing HIV-1 p24 release from CV tissues (Fig. 6).

**Figure 6.**
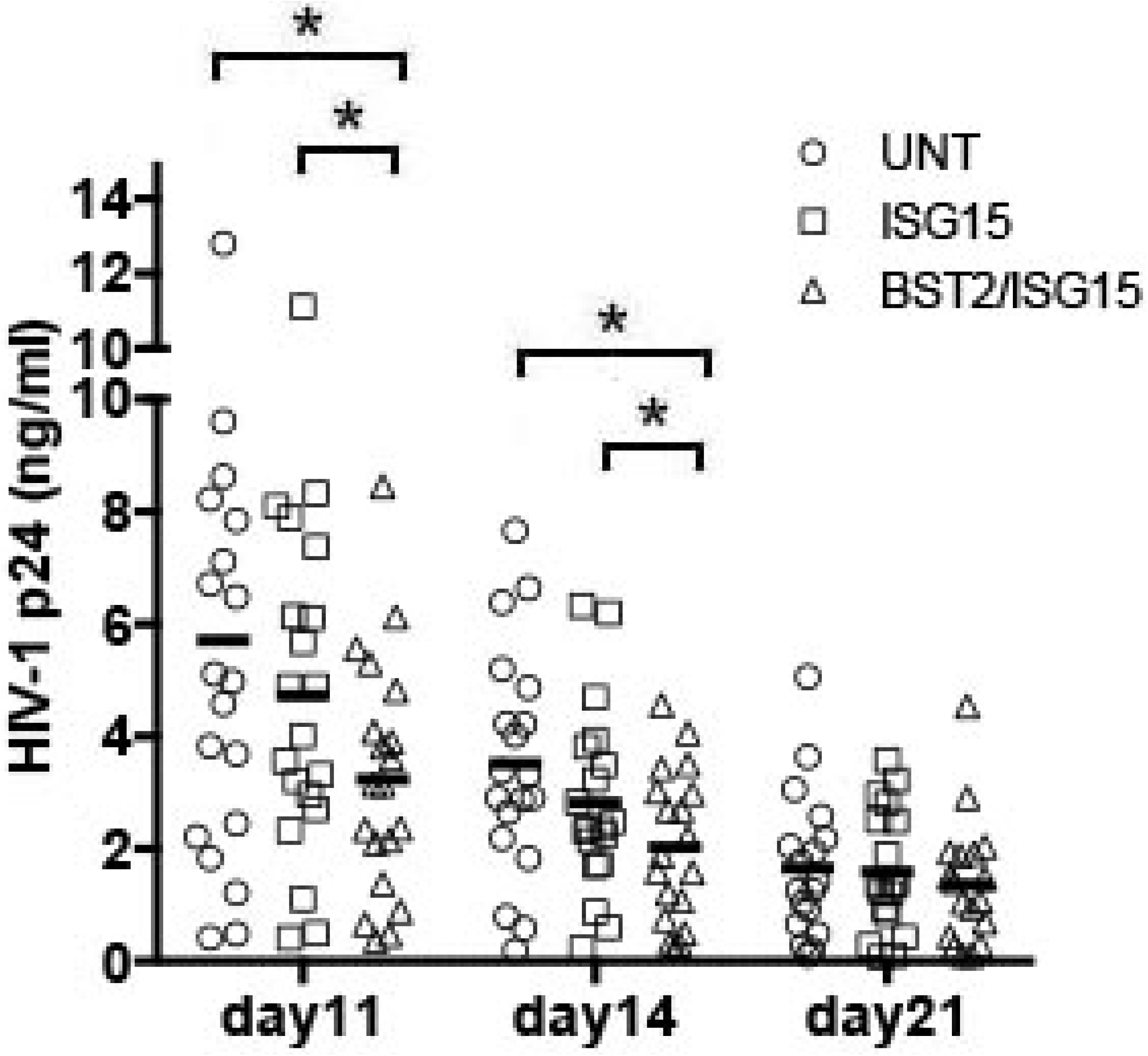
A combination of hrISG15 with hrBST2 is more effective that hrISG15 in reducing HIV-1 replication in CV tissues. HIV-1 p24 levels in CV tissues left untreated or treated with hrISG15 at 0.115 µg/ml alone or in combination with a hrBST2 peptide at 0.1 μg/ml were measured after washing the residual input virus (day 0), and again on days 11, 14 and 21 of infection. Mean values of triplicates from 20 CV tissues are shown. HIV-1 p24 levels on day 0 were below the limit of detection of the assay and are not depicted in the figure* p< 0.05 comparing untreated (UNT) to ISG15/BST2 or ISG15 treated CV tissues.

The decrease in HIV-1 p24 levels was not caused by an hrISG15/BST2 mediated decline in tissue viability as indicated by LDH levels in tissue culture supernatants (data not shown). In contrast to hrIFNα/β, hrISG15/BST2 did not enhance immune cell activation as indicated by similar levels in CD4, CCR5 and CD38 expression between HrISG15/BST2 treated and untreated control tissues (Fig. 7).

**Figure 7.**
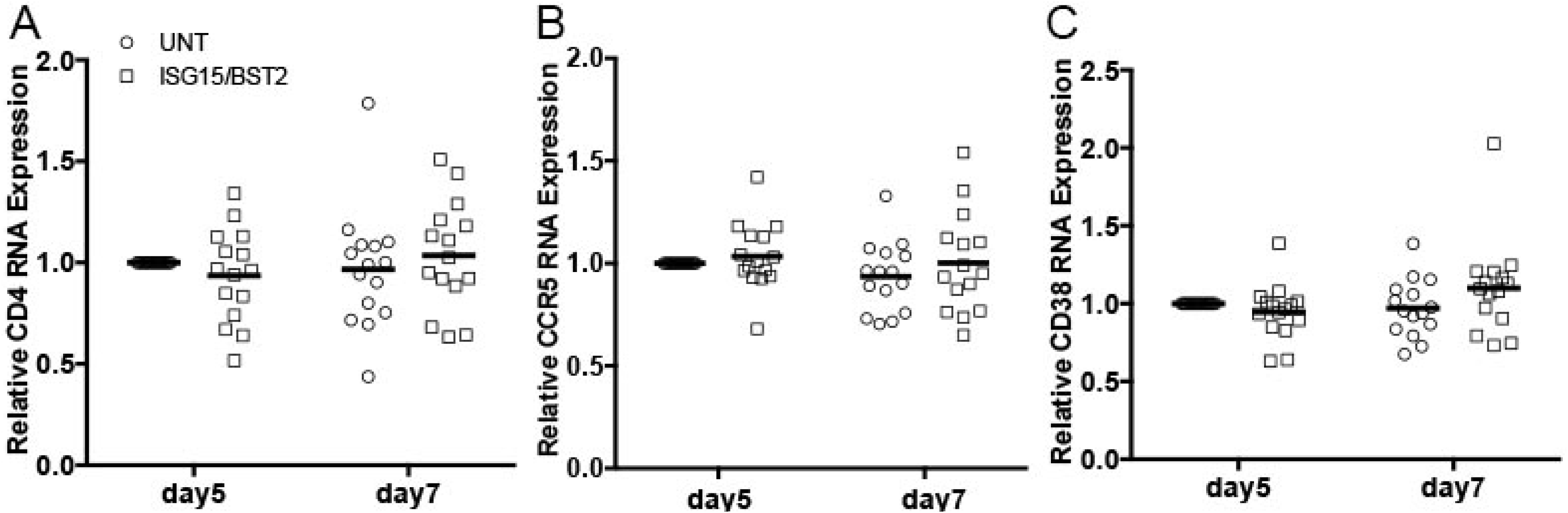
A combination of hrISG15 with hrBST2 does not induce immune cell activation. RNA expression levels of the HIV-1 receptor CD4 (A), co-receptor CCR5 (B) and the immune cell activation marker CD38 (C) in CV tissues left untreated or treated with hrISG15 at 0.115 µg/ml alone or in combination with a hrBST2 peptide at 0.1 μg/ml were quantified by RT-PCR on days 5 and 7 after infection. All data were normalized to GAPDH. For each gene, day 5 values in untreated control tissues were set to 1. Day 5 values in ISG15/BST2 treated tissues or day 7 values in untreated and ISG15/BST2 treated tissues were normalized to day 5 values in untreated tissues. Mean values of triplicates from 15 CV tissues are shown.

## Discussion

Identifying antiviral pathways or target genes with the potential to reduce HIV-1 infection without inducing immune cell activation in the human female genital tract is vital for the prevention of heterosexual HIV-1 transmission in women. This is particularly important in the context of a missing preventative vaccine as well as a partial or no protection of women by the HIV-1 reverse transcriptase inhibitors tenofovir and dapivirine [35-38].

Given that type I IFNs are the first line of defense against sexually transmitted infections, the impact of single hrIFNα or hrIFNβ on SIV, SHIV and HIV-1 replication has been evaluated in *R. macaques*, hu-mice and human primary cells and cell lines [11-19]. Findings on IFNα have been inconsistent, whereas IFNβ reduced HIV-1 infection in these experimental models [11-19,39]. Considering that pathogen binding to immune recognition receptors induces the simultaneous expression of IFNα and β, it is surprising that modulation of HIV-1 infection by both of these antiviral factors has not yet been defined. Consistent with differences in the antiviral activity of IFNα subtypes [40], single IFN α or β applications may not reduce HIV-1 replication compared to the combination in the human CV mucosae, the primary site of viral infection in women [41]. Our manuscript is the first to characterize the effect of combined hrIFNα and β (hrIFNα/β) on HIV-1 replication and immune cell activation in CV tissues. Our findings suggest that hrIFNα/β reduces HIV-1 replication yet enhances immune cell activation in CV tissues. The anti-HIV-1 activity of hrIFNα/β is mediated by stimulation of IFNα/β signaling resulting in enhanced ISG expression. Furthermore, exogenous supplementation with two ISGs induced by hrIFNα/β reduced HIV-1 p24 release to a greater extent than hrIFNα/β without enhancing immune cell activation. Our results support the novel concept that lack of immune cell activation [23] and expression of antiviral genes are key components for decreasing HIV-1 replication in CV tissues. Given that a combination of hrISGs was more effective than hrIFNα/β in reducing HIV-1 p24 release, our findings suggest an advantage of enhancing expression of immune factors downstream rather than activating upstream steps of the IFNα/β signaling pathway.

### HrIFNα/β reduces HIV-1 replication by inducing expression of ISGs in CV tissues

The type I IFN signaling pathway induces expression of both inflammatory and antiviral genes [1,2]. In contrast to single hrIFNα or β, hrIFNα/β reduced HIV-1 p24 release in CV tissues (Fig. 1). This reduction was associated with induction of an antiviral response as indicated by enhanced expression of the IFNAR2 (Fig. 3), the cellular transcription factor IRF7 (Fig. 4B), and the IRF7 induced genes ISG15 and BST2 as well as that of additional ISGs including MX2 and IFITM1 (Fig. 5). These findings demonstrate biological activity of the human recombinant IFN proteins. Given that ISG15, BST2, MX2 and IFITM1 block entry and post-entry steps of the virus life cycle and prevent viral release [2,3,31,32,42-45], our findings support enhancing expression of ISGs as a potential mechanism to reduce HIV-1 replication in CV tissues.

Even though hrIFNα/β reduced HIV-1 replication in the CV mucosae, the antiviral effect was modest and was statistically significant after day 21 of infection. The small reduction in HIV-1 replication is associated with hrIFNα/β enhancing immune cell activation as indicated by up-regulation of CCR5 and CD38 expression (Fig. 2 B-C).

### HrISG combinations decreased HIV-1 replication to a greater extent than hrIFNα/β

Exogenous supplementation with a hrISG15 or a hrBST2 peptide alone did not alter HIV-1 p24 release as indicated by similar HIV-1 p24 levels between tissues treated with single ISG and the untreated control (Fig. 6, and data not shown). Although not statistically significant, CV tissues treated with hrISG15 alone demonstrated a tendency to a decrease in HIV-1 replication. The lack of a significant reduction might reflect high variability in the tissue response to the hrISG15 protein. Since HIV-1 counteracts BST2 [46-48, and our unpublished data], we speculate that tissue supplementation with the hrBST2 peptide alone does not overcome its viral counteraction. Alternatively, BST2 might induce inflammatory factors that neutralize its antiviral effect in CV tissues. Supporting this postulate induction of NFκB by BST2 has been reported [49,50]. In contrast, we found a pronounced reduction in HIV-1 p24 release in tissues treated with a combination of hrISG15 and hrBST2 compared to those exposed to hrISG15 alone or untreated control tissues (Fig. 6). These data suggest a synergistic or additive effect of the ISG combination compared to single ISG. Soluble expression of ISG15 has recently been reported [51, 52]. Given that the molecular weight of the hrISG15 (17 KDa) is above the threshold of 0.9 KDa required for cell penetration our data suggest that the antiviral activity of ISG15 alone or in combination with hrBST2 is due to the soluble molecular form of this protein. HrISG15 in combination with hrBST2 did not enhance CD4, CCR5 and CD38 expression (Fig. 7). Consistent with this finding, exogenous supplementation with hrISG15/BST2 demonstrated early kinetics in reducing HIV-1 replication (day 11) compared to that of hrIFNα/β (day 21) and a more significant decrease (50% reduction). Our findings suggest that activating downstream targets of IFNα/β signaling pathway is a feasible and successful strategy to reduce HIV-1 replication without inducing immune cell activation in CV tissues. Furthermore, an ISG combination is more effective than combined IFNα/β in reducing viral replication.

The mechanisms underlying greater anti-HIV-1 activity of hrISG combination when compared to hrIFNα/β is not understood and beyond the scope of this manuscript. We could speculate that exogenous supplementation with ISGs results in enhanced protein levels when compared to induction of endogenous ISG by hrIFNα/β. Our unpublished data suggest that HIV-1 counteracts the IFNRA2 potentially reducing expression of downstream targets of IFNα/β signaling pathway. Alternatively, hrIFNα/β could induce immune factors that counterbalance the antiviral efficacy of endogenous ISGs. Future experiments will address these questions.

### Mechanisms of interactions between ISG15 and BST2

Innate sensing of HIV-1 by BST2 has been associated with enhanced NFκB-dependent pro-inflammatory responses [49,50]. Given that by stabilizing ubiquitin-specific peptidase 18 (USP18), ISG15 appears to control immune cell activation [51], our findings of a lack of immune cell activation in tissues treated with hrISG15/BST2 (Fig. 7), suggest the potential for ISG15 to control any pro-inflammatory activity of BST2. This effect clearly contributed to the reduced HIV-1 p24 levels observed in hrISG15/BST2 treated tissues.

This study has several limitations. First, we acknowledge data variability as a result of using a relevant human tissue explant model that reflects the variation of women who participate in clinical trials. Our power calculation estimated that we needed to analyze data from at least 15 patients to achieve a 90% power for a type I error of 5 % and an effect size of 90%. All the figures depict findings from either 15 or more patients. By presenting average from triplicates from all of our experiments, we provide the highest possible level of transparency and validate the use of CV tissues to characterize mechanisms of regulation of HIV-1 replication by IFNs and ISGs alone and in combination. Given that we compare experimental treatments and untreated conditions in tissues procured from the same donor, our experimental design controls for all confounding variables and ensures that our findings are indeed due to the treatment. Second, our experimental model does not account for cell recruitment from the peripheral blood to the CV mucosa, thus our findings reflect activation and proliferation of, and the effect of recombinant proteins on, resident immune cells. We speculate that if the experimental model would allow for recruitment of target cells, we would have likely detected a greater antiviral effect of hrISG15 in combination with hrBST2 throughout day 21. It is possible that by not accounting for cell recruitment we might be underestimating the potency of these ISGs to reduce HIV-1 replication.

Third, we only evaluated expression of a subset of ISGs and the anti-viral activity of ISG15 alone and in combination with BST2. Therefore, we are unable to rule out the possibility that additional factors not considered in this study have similar or even greater antiviral effects. Even though we did not test MX2 or IFITM1 for anti-HIV-1 activity in CV tissues several groups have already characterized the mechanisms underlying the antiviral activity of this ISG [42-45]. Enhanced expression of MX2 has been associated with reduced risk to HIV-1 infection in highly exposed seronegative women [53], validating the use of CV tissues to evaluate mucosal ISG expression.

In conclusion, our data suggest that activating downstream targets of the IFNαβ signaling pathway is a feasible and more effective strategy than IFNs treatment to reduce HIV replication without inducing mucosal inflammation. Greater effectiveness could be achieved by ISGs combination compared to supplementation with single ISG.

## Acknowledgments

We thank the Department of Medicine, Section of Anatomic Pathology, DHMC, Lebanon, NH, for their assistance in procuring CV tissues.

## Authors Contributions

Drs. Rollenhagen and Asin participated in the design and execution of the study, interpretation of the results and writing of the manuscript. Dr. Gui conducted the statistical analysis of the data and edited the Statistical Analysis section of the manuscript. Dr. Doncel edited the manuscript.

## References

1. Hoffmann HH, Schneider WM, Rice CM Interferons and viruses: an evolutionary arms race of molecular interactions. TRENDS in Immunology. 2015; 36(3):124–38.

2. Levy DE, Garcia-Sartre A. The virus battles: IFN induction of the antiviral state and mechanisms of viral evasion. Cytokine Growth Factor Rev. 2001; 12(2-3):143–56. Review.

3. Rustagi A, Gale M Jr. Innate antiviral immune signaling, viral evasion and modulation by HIV-1. J Mol Biol. 2014; 426(6):1161–77.

4. Altfeld M, Gale M Jr. Innate immunity against HIV-1 infection. Nat. Immunol. 2015; 16(6): 554–62.

5. Honda K, Taniguchi T. IRFs: master regulators of signaling by Toll-like receptors and cytosolic pattern-recognition receptors. Nat Rev Immunol. 2006; 6(9): 644–58. Review.

6. Honda K, Takaoka A, Taniguchi T. Type I interferon [corrected] gene induction by the interferon regulatory factor family of transcription factors. Immunity. 2006; 25(5):349–60. Review. Erratum in: Immunity. 2006; 25(5):849.

7. Chattopadhyay S, Sen GC. RIG-I-like receptor-induced IRF3 mediated pathway of apoptosis (RIPA): a new antiviral pathway. Protein Cell. 2017; 8(3):165–68. Review.

8. Rollenhagen C, Macura SL, Lathrop MJ, Mackenzie TA, Doncel GF, Asin SN. Enhancing Interferon Regulatory Factor 7 Mediated Antiviral Responses and Decreasing Nuclear Factor Kappa B Expression Limit HIV-1 Replication in Cervical Tissues. PLoS One. 2015; 10(6):1–18.

9. Grandvaux N, Servant MJ, tenOever B, Sen GC, Balachandran S, Barber GN, et al. Transcriptional profiling if IRF3 target genes: direct involvement in the regulation of ISG. J Virol. 2002; 76(11):5532–9.

10. Ning S, Pagano JS, Barber GN. IRF7: activation, regulation, modification and function. Genes Immun. 2011; 12(6):399–414.

11. Sandler NG, Bosinger SE, Estes JD, Zhu RT, Tharp GK, Boritz E, et al. Type I interferon responses in rhesus macaques prevent SIV infection and slow disease progression. Nature. 2014; 511(7511):601–05.

12. Veazey RS, Pilch-Cooper HA, Hope TJ, Alter G, Carias AM, Sips M, et al. Prevention of SHIV transmission by topical IFN-β treatment. Mucosal Immunol. 2016; 9(6):1528–1536.

13. Lavender KJ, Gibbert K, Peterson KE, Van Dis E, Francois S, Woods T, et al. Interferon Alpha Subtype-Specific Suppression of HIV-1 infection In Vivo. J Virol. 2016; 90(13):6001–6013.

14. Abraham S, Choi JG, Ortega NM, Zhang J, Shankar P, Swamy NM. Gene therapy with plasmids encoding IFN-β or IFN-α14 confers log-term resistance to HIV-1 in humanized mice. Oncotarget. 2016; 7(48):78412–78420.

15. Acchioni C, Marsili G, Perrotti E, Remoli AL, Sgarbanti M, Battistini A. Type I IFN--a blunt spear in fighting HIV-1 infection. Cytokine Growth Factor Rev. 2015; 26(2):143–58.

16. Long BR, Stoddart CA. Alpha interferon and HIV infection cause activation of human T cells in NSG-BLT mice. J Virol. 2012; 86(6):3327–36.

17. Goujon C, Malim MH. Characterization of the alpha interferon induced postentry block to HIV-1 infection in primary human macrophages and T cells. J Virol. 2010; 84(18):9254–66.

18. Dianzani F, Castilletti C, Gentile M, Gelderblom HR, Frezza F, Capobianchi MR. Effects of IFN alpha on late stages of HIV-1 replication cycle. Biochimie. 1998; 80(8-9):745–54.

19. Brule F, Khatissian E, Benani A, Bodeaux A, Montagnier L, Piette J, et al. Inhibition of HIV replication: a powerful antiviral strategy by IFN-beta gene delivery in CD4+ cells. Biochem Pharmacol. 2007; 74(6):898–910.

20. Introini A, Vanouille C, Grivel JC, Margolis L. An ex vivo model of HIV-1 infection in Human Lymphoid Tissue and Cervico-vaginal Tissue. Bio Protoc. 2014; 4(4):1–13.

21. Merbah M, Introini A, Fitzgerald W, Grivel JC, Lisco A, Vanpouille C, et al. Cervico-vaginal tissue ex vivo as a model to study early events in HIV-1 infection. Am J Reprod Immunol. 2011; 65(3)268–78.

22. Grivel JC, Margolis L. Use of human tissue explants to study human infectious agents. Nat. Protoc. 2009; 4(2):256–69.

23. Macura SL, Lathrop MJ, Gui J, Doncel GF, Asin SN, Rollenhagen C. Blocking CXCL9 decreases HIV-1 replication and enhances the activity of prophylactic antiretrovirals in human cervical tissues. J Acquir Immune Defic Syndr. 2016; 71(5):474–82.

24. Rollenhagen C, Lathrop MJ, Macura SL, Doncel GF, Asin SN. Herpes simplex virus type-2 stimulates HIV-1 replication in cervical tissues: implications for HIV-1 transmission and efficacy of anti-HIV-1 microbicides. Mucosal Immunol. 2014; 7(5):1165–74.

25. Rollenhagen C, Asin SN. Enhanced HIV-1 replication in ex vivo ectocervical tissues from post-menopausal women correlates with increased inflammatory responses. Mucosal Immunol. 2011; 4(6):671–81.

26. Viswanathan K, Smith MS, Malouli D, Mansouri M, Nelson JA, Früh K. BST2/Tetherin enhances entry of human cytomegalovirus. PLoS Pathog. 2011,7(11):1–13.

27. Warren CJ, Griffin LM, Little AS, Huang IC, Farzan M, Pyeon D. The antiviral restriction factors IFITM1, 2 and 3 do not inhibit infection of human papillomavirus, cytomegalovirus and adenovirus. PLoS One. 2014, 9(5):1–8.

28. Maeda S, Wada H, Naito Y, Nagano H, Simmons S, Kagawa Y, et al. Interferon-? acts on the S/G2/M phases to induce apoptosis in the G1 phase of an IFNAR2-expressing hepatocellular carcinoma cell line. J Biol Chem. 2014, 289(34):23786–95.

29. Palmer CS, Duette GA, Wagner MCE, Henstridge DC, Saleh S, Pereira C, et al. Metabolically active CD4+ T cells expressing Glut1 and OX40 preferentially harbor HIV during in vitro infection. FEBS Lett. 2017; 591(20):3319–3332.

30. Schreiber G. The molecular basis for differential type I interferon signaling. J Biol Chem. 2017; 292(18):7285–7294.

31. Morales D, Lenschow DJ. The antiviral activities of ISG15. J Mol Biol. 2013, 425(24):4995–5008. Review.

32. Okumura A, Lu G, Pitha-Rowe I, Pitha PM. Innate antiviral response targets HIV-1 release by the induction of ubiquitin-like protein ISG15. Proc Natl Acad Sci U S A. 2006;103(5):1440–5.

33. Venkatesh S, Bieniasz PD. Mechanism of HIV-1 virion entrapment by tetherin. PLoS Pathog. 2013, 9(7):1–15.

34. Giese S, Marsh M. Tetherin can restrict cell-free and cell-cell transmission of HIV from primary macrophages to T cells. PLoS Pathog. 2014;10(7):1–21.

35. Abdool Karim Q, Abdool Karim SS, Frohlich JA, Grobler AC, Baxter C, Mansoor LE, et al. Effectiveness and safety of tenofovir gel, an antiretroviral microbicide, for the prevention of HIV infection in women. Science 2010; 329(5996):1168–74.

36. Delany-Moretlwe S, Lombard C, Baron D, Bekker LG, Nkala B, Ahmed K, et al. Tenofovir 1% gel for prevention of HIV-1 infection in women in South Africa (FACTS-001): a phase 3, randomized, double blind, placebo-controlled trial. Lancet Infect Dis. 2018; 18(11):1241–50.

37. Baeten JM, Palanee-Phillips T, Brown ER, Schwartz K, Soto-Torres LE, Govender V, et al. Use of a Vaginal Ring Containing Dapivirine for HIV-1 Prevention in Women. N Engl J Med 2016; 375(22):2121–32.

38. Nel A, van Niekerk N, Kapiga S, Bekker LG, Gama C, Gill K, et al. Safety and Efficacy of a dapivirine vaginal ring for HIV prevention in women. N Engl J Med. 2016; 375(22):2133–43.

39. Schneider WM, Chevillotte MD, Rice CM. Interferon-stimulated genes: a complex web of host defenses. Annu Rev Immunol. 2014,32:513–45.

40. Harper MS, Guo K, Gibbert K, Lee EJ, Dillon SM, Barrett BS, et al. Interferon-α Sub types in an Ex vivo Model of Acute HIV-1 Infection: Expression, Potency and Effector Mechanisms. PLoS Pathog. 2015, 11(11):1–24.

41. Shattock RJ, Haynes BF, Pulendran B, Flores J, Esparza J; Working Group convened by the Global HIV Vaccine Enterprise. Improving defences at the portal of HIV entry: mucosal and innate immunity. PLoS Med. 2008, 5(4):537–541.

42. Kane M, Yadav SS, Bitzegeio J, Kutluay SB, Zang T, Wilson SJ, et al. MX2 is an interferon-induced inhibitor of HIV-1 infection. Nature. 2013, 502(7472):563–6.

43. Goujon C, Moncorgé O, Bauby H, Doyle T, Ward CC, Schaller, et al. Human MX2 is an interferon-induced post-entry inhibitor of HIV-1 infection. Nature. 2013, 502(7472):559–62.

44. Lu J, Pan Q, Rong L, He W, Liu SL, Liang C. The IFITM proteins inhibit HIV-1 infection. J Virol. 2011;85(5):2126–37.

45. Yu J, Liu SL. The Inhibition of HIV-1 Entry Imposed by Interferon Inducible Transmembrane Proteins Is Independent of Co-receptor Usage. Viruses. 2018;10(8):1–16

46. Kmiec D, Iyer SS, Stürzel CM, Sauter D, Hahn BH, Kirchhoff F. Vpu-Mediated Counteraction of Tetherin Is a Major Determinant of HIV-1 Interferon Resistance. MBio. 2016;7(4):1–10.

47. Schindler M, Rajan D, Banning C, Wimmer P, Koppensteiner H, Iwanski A, et al. Vpu serine 52 dependent counteraction of tetherin is required for HIV-1 replication in macrophages, but not in ex vivo human lymphoid tissue. Retrovirology. 2010; 7(1):1–13.

48. Mangeat B, Gers-Huber G, Lehmann M, Zufferey M, Luban J, Piguet V. HIV-1 Vpu neutralizes the antiviral factor Tetherin /BST-2 by binding it and directing its beta-TrCP2-dependent degradation. PLoS Pathog. 2009;5(9):1–12.

49. Galão RP, Le Tortorec A, Pickering S, Kueck T, Neil SJ. Innate sensing of HIV-1 assembly by Tetherin induces NFкB-dependent proinflammatory responses. Cell Host Microbe. 2012; 12(5): 633–44.

50. Tokarev A, Suarez M, Kwan W, Fitzpatrick K, Singh R, Guatelli J. Stimulation of NFкB activity by the HIV restriction factor BST2. J Virol. 2013;87(4):2046–57.

51. Hermann M, Bogunovic D. ISG15: In Sickness and in Health. Trends Immunol. 2017; 38(2): 79–93. Review.

52. Swaim CD, Canadeo LA, Huibregtse JM. Approaches for investigating the extracellular signaling function of ISG15. Methods Enzymol. 2019; 618:211–227.

53. Stein DR, Shaw SY, McKinnon LR, Abou M, McCorrister SJ,Westmacott GR, et al. MX2 expression is associated with reduced susceptibility to HIV infection in highly exposed HIV seronegative Kenyan women. AIDS. 2015, 29(1):35–41.

